# Longitudinal brain studies in adult zebrafish by MRI

**DOI:** 10.1101/2022.06.09.495545

**Authors:** Noémie Hamilton, Claire Allen, Steven Reynolds

## Abstract

Zebrafish (Danio rerio) has been successfully used for decades in developmental studies and disease modelling. The remarkable intake of zebrafish as a model system is partly due to its transparency during the early weeks of its development, allowing *in vivo* imaging of cellular and molecular processes. However, this key advantage wears off when tissues become opaque as the animal reaches juvenile and adult stages, rendering access to tissues for live imaging and longitudinal studies difficult. Here we aimed to provide a novel approach to image and assess tissue integrity of adult zebrafish using Magnetic Resonance Imaging (MRI) on live zebrafish suitable for longitudinal studies. We built a 3D-printed life support chamber and designed a protocol-directed sedation regime to recover adult zebrafish post scanning in a 9.4T MRI scanner. Our life support chamber is cheap and easy to create using 3D printing, allowing other groups to copy our template for quick setup. Additionally, we optimised the delivery of contrast agent to enhance brain signals in order to refine current delivery, usually delivered by intravenous in rodents. We show here that immersion in gadolinium was a viable alternative to intraperitoneal injection to reduce T1 relaxation times. This could lead to improved image contrast in adult zebrafish disease models. In conclusion, we provide here a detailed methodology to allow longitudinal studies of brain tissue integrity of adult zebrafish, combining safe and efficient delivery of contrast agent and live MRI. This technique can be used to bridge the gap between *in vivo* studies and longitudinal brain analysis in adult zebrafish which can be applied to the ever-growing number of adult zebrafish models of ageing and neurodegenerative diseases.

## Introduction

Neurodegenerative diseases are devastating and affect millions of people in the UK. Animal models are essential in characterising the disease in order to understand brain tissue morphology, integrity and function. In accordance with the refinement principle of the 3Rs (replace, reduce, refine), neurodegenerative research has steadily transitioned from rodent to zebrafish models, which are a lower neurophysiological, versatile and cost-effective species. Zebrafish are a popular research model for studying neurological diseases, such as Parkinson’s, multiple sclerosis, leukodystrophies and epilepsy (Burrows et al., 2019; Hamilton et al., 2020; Keatinge et al., 2015; Najib et al., 2020; Rutherford & Hamilton, 2019; Yaksi et al., 2021). Most of the work into these neurological conditions is conducted on transparent juvenile fish, which are amenable to optical imaging techniques. However, longitudinal monitoring of disease progression is visually limited by the opacity of adult zebrafish restricting the scope of data collection at these later stages.

Magnetic Resonance Imaging (MRI) is a well-known non-invasive medical tool for diagnosing patients suffering from neurodegenerative diseases and is ideally suited for longitudinal studies. It can image both tissue integrity and function throughout the brain with rodent models underpinning much of this work (Denic et al., 2011; Dijkhuizen & Nicolay, 2003). Preclinical MRI has previously been used to study disease models of zebrafish, where the application of ultrahigh field (>7T), powerful gradient systems and bespoke small volume coils permits relatively high-resolution images - on the order of 10s μm. Most of these have used fixed tissue fish (Haud et al., 2011; Ullmann et al., 2009), however, more recently publications reported MRI scanning of live fish (Kabli et al., 2009; Koth et al., 2017; Merrifield et al., 2017). Life support for the fish has required the implementation of an MRI compatible flow tank, which can produce turbulence leading to a degradation in imaging quality. Furthermore, a wider field of view (FOV) is required that encompasses the flow chamber to prevent wrap around effects and necessitates more phase encoding steps, i.e. longer acquisition times, for a given resolution.

We have developed a prototype life support chamber for zebrafish that removes the need for a water filled flow chamber. The chamber can be used in a vertical bore MRI scanner with anaesthesia supplied by readily available laboratory equipment, i.e. syringe pumps. Supplying of anaesthesia by intubation directly to the fish mouth removes the need for water to surround the gills, therefore reducing turbulence as the water drains down the flanks of the fish keeping it moist, and minimises fish movement. Additionally, the FOV in the phase encoding direction can be reduced by placing it along the sagittal axis of the fish. Cost and ease of fabrication were prioritised during the design of the chamber: consideration was given to availability of parts and materials, the chamber is 3D printable, which is cheap and rapid, allowing the chamber either to be reused or could be a single-use item if, for instance, spread of infection was a concern. The live chamber permitted scanning of anaesthetised adult zebrafish for 2 hours, followed by full recovery.

Gadolinium based contrast agents (GCA) are commonly used as a method to increase tissue contrast and/or reduced imaging time (Wahsner et al., 2019). These agents have also been applied to zebrafish MRI for live imaging in the heart (Koth et al., 2017) and fixed or freshly sacrificed fish brain tissue (Ullmann et al., 2009). Whilst intra peritoneal (IP) injection is a commonly accepted administration route for small animals it can lead to tissue injury or mis-dosing due to the small volumes of injected fluid. In this study, we allowed fish to swim freely immersed in GCA mixed to fish water to examine its effect on T1 and T2 values and compared this to IP injected animals. Our study has developed a novel chamber to safely hold a zebrafish for MRI and refined current contrast agent administration protocol to comply with the 3Rs approach.

## Results

### 3D-printed life support chamber combined with dual anaesthetic regime allows acquisition of MR images in live zebrafish with full subsequent recovery

In order to image a live adult zebrafish for a prolonged period of time, in a safe, consistent and reproducible way, we developed a life support chamber to sustain water to the animal (Figure 1). This kept the fish out of water to reduce imaging artifacts from water passage. Adult zebrafish were held vertically using four flexible rods that can adapt to different body sizes and shapes without crushing the fish. The rods are transparent and have an open design which helps with the insertion of the fish and final positioning (Figure 1A). Oxygenated water containing a sedative was delivered with a constant flow using a dual syringe pump through a thin PVC tube that was guided from the top of the chamber directly into the fish mouth (Figure 1B). To keep the intubated fish hydrated during image acquisition, the water expelled by the fish gills was caught by a soft cloth wrapped around its body and a sponge placed under the tail (Figure 1A). To prevent water leakage into the MRI machine the fish and tubing were enclosed in a glass tube, sealed with O-rings and water collected at the bottom was evacuated by a waste tube (Figure 1A). Once the fish was intubated, positioned and sealed under the glass tube, the chamber was inserted and secured into the MRI probe then inserted into the scanner. The whole process took only a few minutes. A step-by-step protocol is provided as a supplemental file.

**Figure 1:**
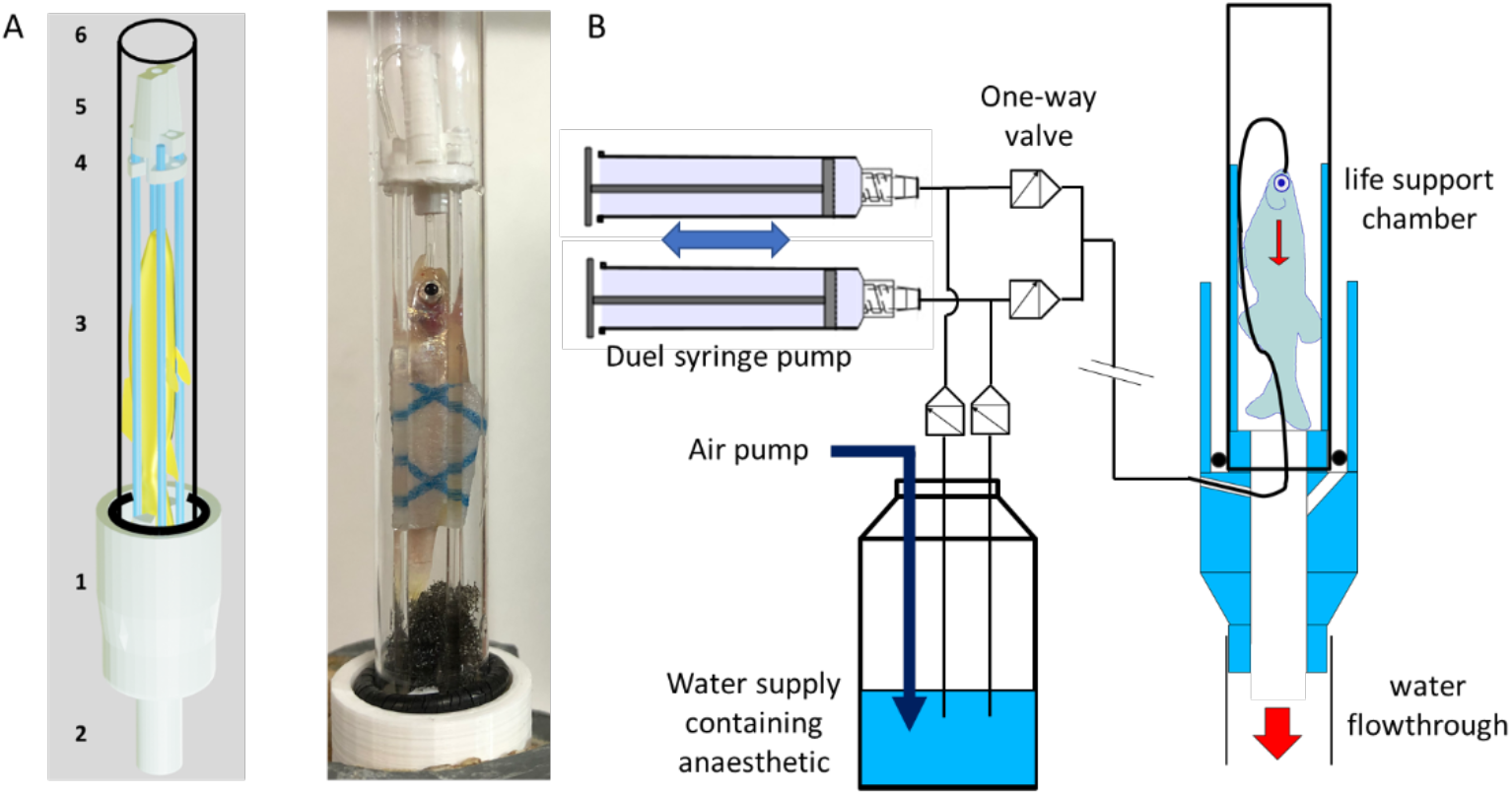
3D-printed chamber supports zebrafish adults during MRI scans. A) The fish life support chamber composed: main body (**1**) with water for drainage (**2**), four flexible fish support rods (**3**) are held by a retaining plate (**4**). Water supply passed into the fish’s mouth via a guide (**5**). Water tightness was made by a 10 mm glass NMR tube (**6**) that fits over the upper part of the assembly and is sealed to the main body (**1**) with o-rings. B) A constant flow water supply system was provided to the fish by two syringe pumps. Oxygenated water was drawn from a main reservoir that contained anaesthetic (see main text). The pumps worked in push/pull tandem so that one is filling as the other supplies water to the fish. See main text for details.

We implemented an anaesthetic regime to minimise stress and allow full recovery of the animal post scan. Tricaine methanesulfonate (MS-222) is the most common anaesthetic used in zebrafish embryos as it immobilised animals rapidly and efficiently for up to 48h (Kaufmann et al., 2012; Kimmel et al., 1995; Westerfield, 1995). However, MS-222 induces cardiac arrest in adult zebrafish after exposure as short as 15 minutes, therefore rendering its use limited for long term imaging and longitudinal studies (Huang et al., 2010; Monte & Zoltán, 2012). A combination with either the compound Benzocaine or Isoflurane was found to decrease risks of cardiac arrest and improve recovery (Huang et al. 2010; Lockwood et al. 2017; Wynd et al. 2017). Benzocaine is also known to allow longer staged anaesthesia (Olt et al., 2016). We therefore designed a dual anaesthetic regime using both MS-222 and Benzocaine to fine-tune the depth of anaesthesia during the entire process (Figure 2). MS-222 at 160mg/ml was used to initially sedate the fish and keep immobilised while handling and wrapping in cloth. The fish was then moved into the chamber, intubated and positioned while receiving MS-222 at 120mg/ml through a drip on the head and through intubation. Once inserted into the probe, the anaesthetic was switched to Benzocaine at 35mg/L for the duration of the scan (up to 2h). For this project, we scanned a total of 29 fish and all were fully recovered but 2 due to an overdose from MS-222.

**Figure 2:**
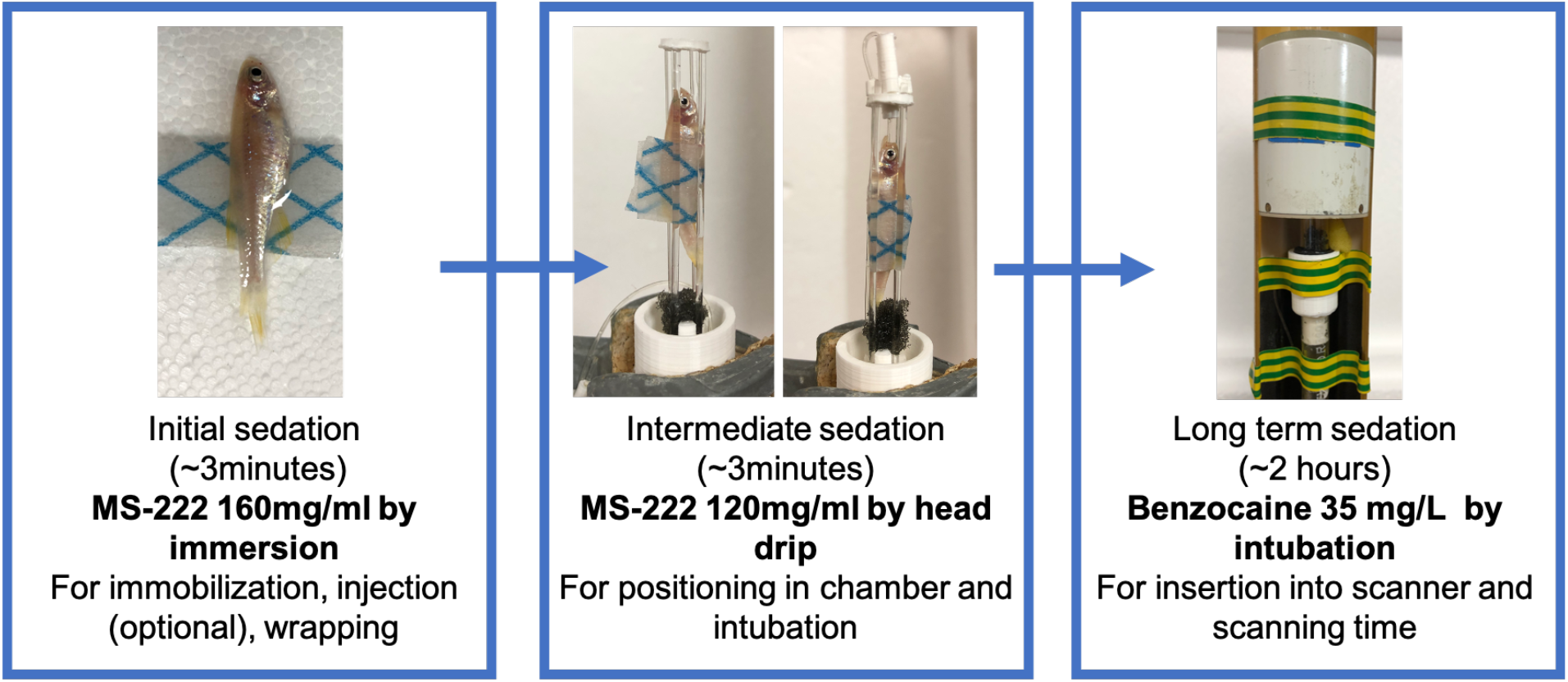
MS-222/Benzocaine dual anaesthetic regime allows recovery of zebrafish after 2h MRI scans.

Using the life support chamber combined with the anaesthetic regime, we then tested the efficiency of the system to produce MRI scans clear of motion artefacts, such as water passage. Using MSME (Multi-Slice Multi-Echo) or RARE (Rapid Acquisition with Refocused Echoes) scans, images were obtained at 50 μm in plane resolution and 0.2 – 0.5 mm slice thickness, requiring 30 – 90 min to acquire depending on slice thickness and number of averages (Figure 3 and gif video Figure S1). By calculating of the Signal to Noise Ratio (SNR), we showed that images of live fish were comparable to those from fixed fish, with SNR higher in live fish compared to fixed fish, although this was not significant by a paired Wilcoxon Signed Rank Test, see Table 1.

**Figure 3:**
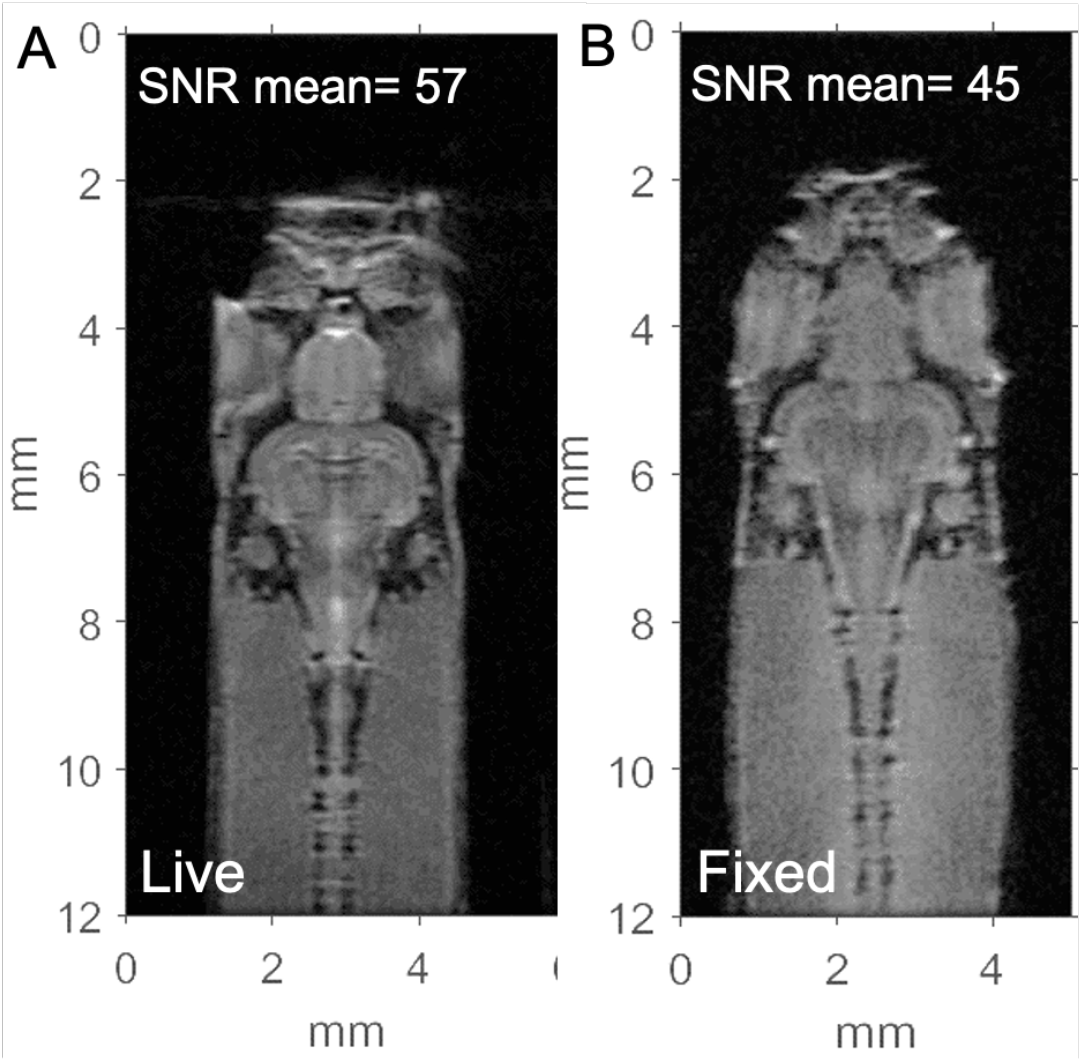
Signal to noise (SNR) is similar between live and fixed scans. Representative spin echo images (RARE) from the same live (**A**) then fixed (**B**) zebrafish, with 50×50mm in plane resolution, 200mm slice thickness, TE/TR 14/1500ms, NEX 64. Signal to noise (SNR) mean across 3 animals calculated within the brain region of interest. **A**. FOV 12×6mm 96 min. **B**. FOV 12×5mm, 80 min.

**Table 1:** Signal to Noise Ratio (SNR) for live and fixed fish within the brain ROI.

| Treatment | N | SNR, mean (range) |  |  |  |
| --- | --- | --- | --- | --- | --- |
|  |  | Live | Fixed | Ratio<sup>‡</sup> | P-value<sup>†</sup> |
| GCA, injected | 3 | 59 (55 – 67) | 45 (37 – 50) | 1.34 | 0.25 |
| GCA, immersion | 5 | 70 (47 – 84) | 52 (40 – 60) | 1.41 | 0.19 |
| PBS, injected | 3 | 65 (60 – 70) | 47 (40 – 53) | 1.40 | 0.25 |
| control | 3 | 57 (51 – 63) | 45 (41 – 51) | 1.28 | 0.25 |

<sup>‡</sup>SNR<sub>live</sub>/SNR<sub>fixed</sub> in the same animal (all brain ROI image slices), with the mean calculated per treatment
<sup>†</sup>Wilcoxon sign rank test

### Immersion in contrast agent refines common injection procedure for brain integrity analysis

To enhance SNR and reduce scanning time, we tested the addition of Gadolinium contrast agent (GCA) using the common Intra Peritoneal (IP) injection alongside a less invasive immersion approach. For each treatment group, heatmaps of T1 and T2 measurement were generated for each slice containing brain tissue (Figure 4).

**Figure 4:**
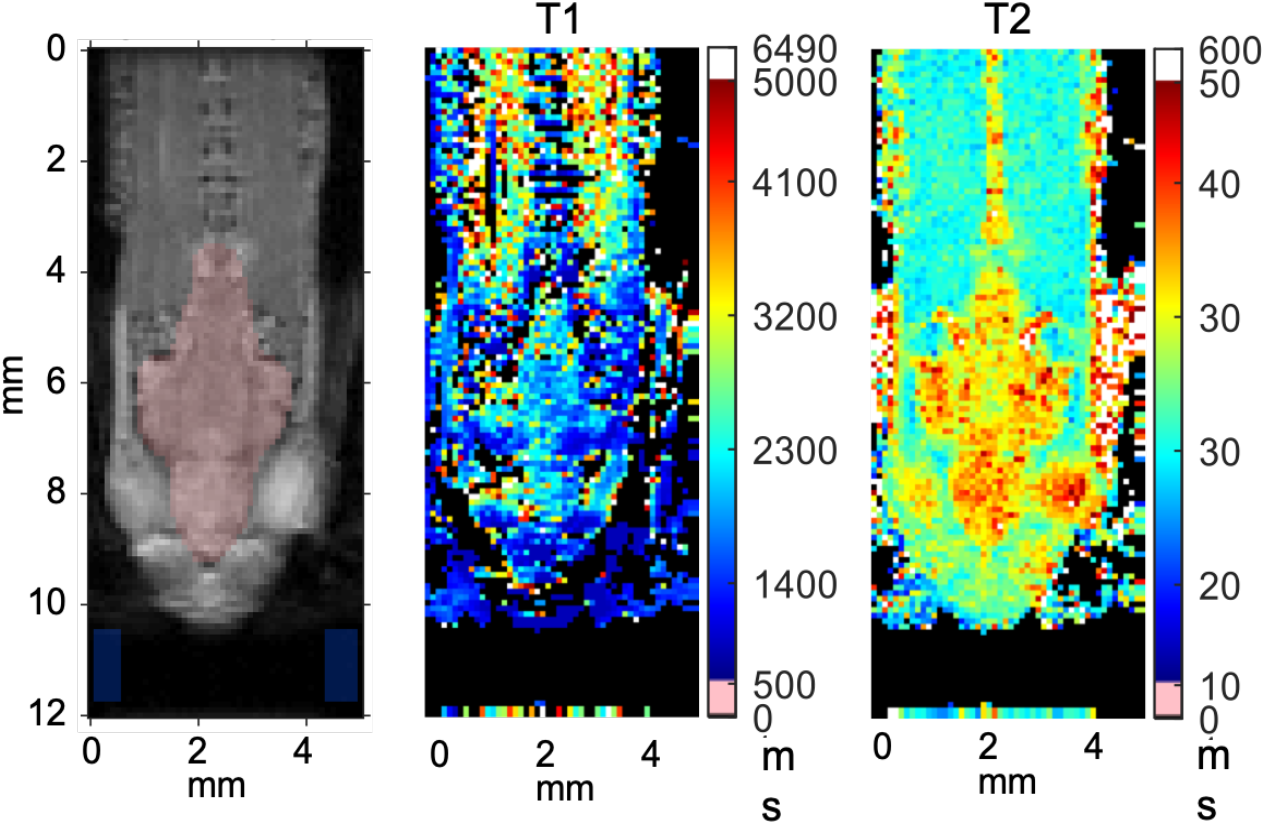
Brain region of interest (ROI) used for T1 and T2 measurement. Example of RARE-VTR images showing magnitude (left, TE = 10 ms, TR 10000 s); T1 map (middle); T2 map (right); brain ROI in red. In the bottom left and right corners of the magnitude image can be seen the location of noise regions in each slice used to calculate the signal to noise ratio (see methods for details). The colour bars to the right of the T1 and T2 maps depict the range of values. For clarity, the colour range was restricted such that values greater than 5000 ms (T1) or 50 ms (T2) are all set to white and values less than 500 ms (T1) or 10 ms (T2) are set to pink.

As expected, administration of GCA by injection caused a significant reduction in T1 values of brain tissue compared to control fish groups (PBS injected and untreated; Figure 5A), mean T1 values (1590 ± 27 ms) that were significantly lower than all other groups. The immersion method reduced T1 values (2341 ± 20 ms) significantly lower than control groups: PBS injected (2955 ± 20 ms) and untreated fish (3426 ± 19 ms) (Figure 5A and Table S1 for p-values). Two animals from the GCA immersion group had T1 values that were not significantly different from control fish, with one of them containing motion artefact (Figure S2 and S3). Nevertheless, immersion in GCA still provided an imaging advantage compared to the control groups and can be used to improve image quality instead of the more invasive IP injection (Figure 5A).

**Figure 5:**
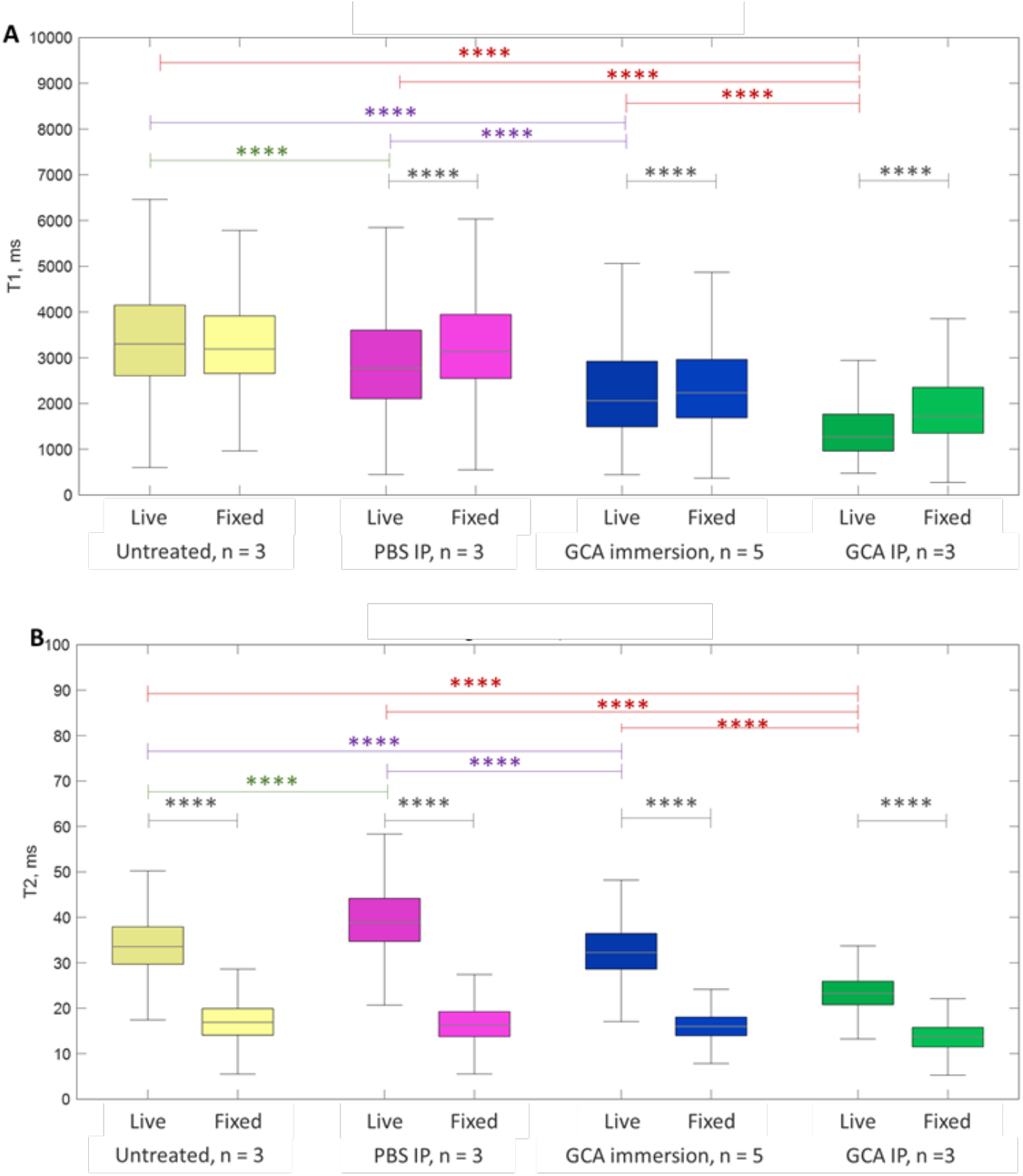
Gadolinium contrast agent (GCA) immersion improves image acquisition. T1 (A) and T2 (B)-relaxation time measurement within the brain ROI of both live and fixed zebrafish. Prior to scanning, live fish were treated with GCA by IP injection or immersion, PBS by IP injection or left untreated. The boxplots show median and interquartile range (IQR), with whisker located at 1.5*IQR. Outliers have been removed for clarity and y-axis scaling (The full plot can be view in supplementary Figures S3 and S4 and outliers were included in the statistical analysis). The number of outliers represented 0.2 – 10.4% of the total number of voxels. Horizontal bars show Bonferroni Post-hoc differences between live fish. Statistical comparisons between the fixed fish groups are not shown. See supplementary Tables S1 and S2 for live and fixed fish details respectively. ****, p << 0.0001.

As per T1 images, we observed significant changes in T2 values in the GCA treated versus PBS/control fish (Figure 5B and supplementary Table S2 for p-values). Although IP injection of GCA showed the lowest T2 values (24.3 ± 0.2 ms), GCA immersion values (32.8 ± 0.1 ms) were still significantly lower than control (34.4 ± 0.1 ms) and PBS injected (40.4 ± 0.2 ms) groups. Of note, T2 values for three of the GCA immersion fish were not significantly different from PBS injected/control fish, two of these were the same animals that also did not show significant change for the T1 data (Figure S5 and S6).

To provide a comparison to fixed tissue MRI and assess GCA effects this tissue, each animal scanned live were subsequently culled by immersion fixation in 4% PFA and scanned again 48h post fix. Overall, and irrespective of the treatment with GCA, the brain tissue of fixed fish showed T1 values that were significantly higher than in live fish for all groups except free swimming controls (Figure 5A and Table S3). Fixed fish T1 values for GCA treatment followed the same trend as for the live fish, with the exception that T1 values in PBS injected which were not significantly different from control animals (Table S1). T2 values were very similar between all fixed groups with significant difference only in T2 values for GCA treated groups (Tables S2, S3). T2 values were significantly higher for the live brains compared to fixed samples (Figure 5B and Table S3).

We then asked whether T1 and T2 values could be extracted from different brain regions to provide a more detailed analysis. ROIs were manually drawn to separate the forebrain, the midbrain and the hindbrain region (Figure S7). In live fish there were no significant differences in T1 values between the ROIs (fore, mid or hind) irrespective of whether gadolinium had been administered or not (Figure S8a. However, after the fish had been fixed, T1 values were significantly different between brain ROIs for GCA immersion, PBS injected and control fish (Figure S8b and Table S4 for T1 and p-values). In these three fixed fish cohorts, the fore brain ROI was significantly higher than the mid and hind ROIs. In contrast, there were no significant differences were observed between brain ROIs for fixed fish injected with GCA.

In the case of brain ROI separated T2 values significant differences in T2 values were observed for live fish, see supplementary Figure S9a. For each cohort, T2 values were highest in the fore brain region of GCA immersion, PBS injected and control fish, whilst the mid-brain ROI showed the highest T2 values for GCA injected fish (Table S5). A similar observation in the brain ROIs was made for fixed fish T2 values with significant difference between forebrain and midbrain ROI in GCA injected, GCA immersion and PBS injected fish (as well as between fore and hind ROIS in the latter two cohorts). No significant differences were observed in the control group (Figure S9b and Table S6).

## Discussion

We describe here the first vertical live support chamber and its use for MRI scanning of adult zebrafish which we used to refine current injection protocol of contrast agent by using non-invasive immersion instead. Recently, a few bespoke chambers have been developed to sustain a live zebrafish during MRI scanning. These contain fully or partial water filled chambers (Kabli et al., 2009; Koth et al., 2017; Merrifield et al., 2017), whereas the chamber presented here used a more open design. This allowed the animal to be observed for well-being prior to insertion into the magnet. Despite placing the fish in a vertical position for more than an hour there were no obvious signs of stress to the animal and the vast majority (27 out of 29) recovered within a few minutes when returned to freshwater. The open chamber design was also advantageous in that fish could be rapidly exchanged, freely positioned vertically depending on their size. This versatility allowed the fish to be located such that the anatomical region of interest could be placed at the most sensitive region of the radio frequency (rf) coil. In principle, the fish could be moved sequentially in the vertical to image other anatomical regions. The chamber could easily be adapted to accept fish of varying sizes and could be redesigned for larger species that fit within the volume of rf coils that were available to a facility.

The imaging resolution obtained in this study was comparable to those previously described (Koth et al., 2017; Merrifield et al., 2017). Generally, the image quality for live fish was similar to that of fixed fish, with a 30 – 40% improvement in SNR for live fish – most likely due to tissue fixative degrading the signal. The available signal to noise and imaging time ultimately limits the practical resolution and slice thickness that can be used, without excessively long anaesthetic times for the animal, due to signal averaging. On the other hand, fixed fish can be scanned for as long as one desires to achieve the required image resolution and SNR performance, but with the obvious lack of options for continuation of longitudinal studies. We limited our fish to less than 2 hours of sedation. Improvements to rf coil design would help here, particularly if cryo-cooled coils could be developed. Alternatively, compressed sensing and systemic administration of gadolinium-based contrast agents could reduce scanning time due to T1 shortening, further reducing anaesthesia time.

T1 values were higher in the control fish in this study (control live fish T1, 3426 ± 19 ms) compared to those previously reported for freshly sacrifice fish (T1 1000 – 2000 ms; T1 2593 ± 138 ms, respectively) (Kline et al., 2019; Ullmann et al., 2009). The reason for the lower reported values by the Kline study could be due to experiments being performed at 4°C and 7T. The temperature dependence of T1 has been shown in post-mortem human brain tissue, where T1 values can change by 3 - 17 ms/°C (T2 changes were more modest. -0.2 – 0.3 ms/°C)(Birkl et al., 2016). Additionally, the T1 values increase with magnetic field strength. Ulman *et al* estimated T1 values at a higher field strength (16.4T) than ours (Ullmann et al., 2009). However, they reported values from a single mid-sagittal slice (0.1 mm slice thickness) containing the telencephalon, hypothalamus and cerebellum, compared to our values for whole brain. Although T1 values were different, the T2 values in our study (control live fish T2, 34.1 ± 0.1 ms) were comparable to those from the Kline *et al* and Ullman *et al* studies (freshly sacrificed, T2, 30 – 50 ms; 22.13 ± 2.2 ms respectively). At low field clinically relevant magnetic field strengths T2 relaxation remains largely constant. At ultrahigh field, >7T, microscopic diffusion and magnetic susceptibility gradients can lead to a reduction in T2 values (de Graaf et al., 2006).

The type of fixative applied to brain tissue has been reported to effect both T1 and T2 values in zebrafish (Ullmann et al., 2009). Unlike Ullman et al we found a significant reduction in T2 values for fixed untreated fish compared to untreated live fish. This may have been due to the longer fixative immersion times used by us (>12h) compared to Ullman (3.5h) and similar T2 reduction for longer fixation time were described by Thelwall et al, who applied fixative over a number of weeks (Thelwall et al., 2006; Ullmann et al., 2009). T2 relaxation is sensitive to molecular motion and the addition of fixative changes the physical properties of the tissue, as well as potentially changing the exchange of protons between water molecules (Ullmann et al., 2009). This influences T2 more than T1 – which is driven mainly by intramolecular dipolar relaxation between nuclei, well as with unpaired electrons present in a contrast agent. The significant variation in T1 and T2 values with brain region (fore, mid and hind) may be in part due to partial volume effects given the relatively large voxel size (100 μm) and slice thickness (500 μm). However, the similarity in T1/T2 values between our and other studies should result in similar image contrast and quality for a given MRI imaging protocol.

Gadolinium-based contrast agents have been used before in live zebrafish to shorten scanning time of the heart (Koth et al., 2017), but to our knowledge T1 and T2 values have only been reported in sacrificed fish (Kline et al., 2019). We investigated whether immersion of the fish in GCA doped tank water could be a viable alternative administration route to intra-peritoneal (I.P) injection. Immersion administration of GCA offers an alternative to I.P. injection. The reduction in T1 and T2 values were between those of the I.P. administered GCA and the control cohorts (PBS, I.P. and control). However, in two of the five experiments there was no significant change in T1 or T2 values for the GCA immersion experiment. These experiments were performed as the others, and it is unclear as to the reason why the fish did not take up the GCA in sufficient quantities. However, GCA immersion should be considered where repeated IP injection could damage tissue, technical expertise in IP injection is not available or the small volume used can lead to IP dosing variation. In our study, the GCA concentration and immersion time was not optimised. The immersion time was taken for pragmatic reasons whilst one of the other fish treatment groups was scanned. It is possible that similar changes T1/T2 could be achieved with shorter immersion times (<2h). Additionally, modifying the concentration of GCA could achieve a greater effect. However, it should be noted that the fish required a minimum of 100 ml tank water to swim in and necessitates the use of several millilitres of GCA for a specific concentration, whereas IP injection only requires a few microlitres.

By providing longitudinal analysis of zebrafish brain tissue in later life, we hope that MRI will help to widen the use of zebrafish in modelling neurodegenerative diseases and its progression.

## Materials and Methods

### Zebrafish husbandry and ethics

All zebrafish were raised in the Biological Services Aquarium at the University of Sheffield in the UK Home Office approved aquarium and maintained following standard protocols(Nüsslein-Volhard & Dham, 2002). Tanks were maintained at 28°C with a continuous recirculating water supply and a daily light/dark cycle of 14/10 hours. All procedures were performed on adult zebrafish to standards set under AWERB (Animal Welfare and Ethical Review Bodies) and UK Home Office–approved protocols (Project Licence - P254848FD). Fish were transported to the MRI facility in a polystyrene carrier box containing a single tank. Fish were kept in the box to keep a constant temperature in a designated animal room until needed for experiment on the same day. The MRI facility suite is kept air conditioned at 21°C and no heating was provided to the fish whilst they were in the scanner.

### Zebrafish life support chamber

The fish life support chamber was intended to supply water directly to the animal without the exterior water surrounding the fish. It was designed to be inserted into a 10 mm MRI volume coil and provide easy access for fish loading and observation. The chamber was modelled in open-source software Blender (www.blender.org) and 3D printed using Cura software and an Ultimaker 2+ Extended 3D printer (both Ultimaker, Utrecht, Netherlands). Objects were printed with polylactic acid (PLA), see Figure 1. The PLA main body (1 in Figure 1) supports all components and collects water for drainage (2). The zebrafish was supported (3) by four flexible acrylic rods (1 mm diameter, ∼5 cm in length. <u>4D Modelshop Ltd</u>, London, UK) that slot into the main body and a PLA retaining plate (4) at the top. Water was supplied to the fish via a PVC tube (not shown on the 3D model) that passed into the side of the main body, up past the fish, through a PLA guide (5) and into the fish’s mouth. The compartment was made water tight by a 10 mm glass NMR tube (6) that fitted over the upper part of the assembly and was sealed to the main body (1) with an o-ring. To maintain the fish physiology, oxygenated water (containing anaesthetic, see below) passed into the fish’s mouth, out via the gills and down their flanks. Water was drained through a long tube to the bottom of the probe and into a collecting jar. To allow for adjustment of the water supply tube it was not sealed to the main body but relied on the design of the chamber, i.e. the hole slope downwards into the chamber, so it could be adjusted for the fish, whilst preventing water egress through its access hole.

### Zebrafish Water supply

The tube inserted into the fish’s mouth consisted of a short, ∼20 cm, 0.25 mm I.D tube that was extended with a 0.89 mm I.D tube approximately 2 m in length (S3 and S54-HL Tygon respectively, Cole Parmer, St Neots, UK). Continuous water supply was maintained by two Aladdin NE-1000 syringe pumps (World Precision Instruments, Hitchin, UK) that were synchronised to draw oxygenated water from a main reservoir that contains anaesthetic. The pumps were programmed to work in tandem such that one filled as the other supplied water to the fish at 3 ml/min. 10 ml syringes were used so that each one cycles over ∼5 minutes (fill/discharge).

### Zebrafish anaesthesia, scanning and recovery

Tricaine/MS222 (Sigma – Cat. No. E10521, Gillingham, UK) stock was made at 4 g/L in filtered system water, with pH adjusted to 7.4 using 1 M Tris. A brown glass bottle protected the stock solution from light and was stored in the fridge. Subsequent tricaine solutions were made using Tricaine stock and E3 water (10x E3: 5 mM NaCl, 0.17 mM KCl, 0.33 mM CaCl2, 0.33 mM MgSO4 diluted to 1x E3 with distilled water (Westerfield, 1995). The anaesthetic solution concentrations were made with high accuracy: 160 mg/ml of tricaine (4ml in 96ml of E3) and 120 mg/ml tricaine (3 ml in 96 ml of E3). Benzocaine (Sigma E1501) stock was made at 50 mg/ml by dissolving in 70% ethanol. This was aliquoted into 750 μl and stored at -20°C until usage. Benzocaine was used at the final concentration of 35 mg/L (using 700 μl in 1 L to directly feed the syringe pumps).

For initial anaesthesia, fish were transferred using a net to a beaker containing 100 ml of 160 mg/ml tricaine. When the fish stopped moving and responding to touch, it was lifted out of the beaker and transferred to a shallow bath of 160 mg/ml tricaine in a clean petri dish. There, the fish was wrapped into a thin rectangle of multipurpose cloth (Spontex CODE/SKU: 19900074), which allowed the fish to be gently held vertically without slipping or damaging its scales. Fish were vertically inserted between the pliable rods of the live chamber (see above), ensuring the feeding tube was positioned ventrally to stop any water artefacts appearing in the image. During the positioning process, the fish was kept moist by dripping drops of 160 mg/ml tricaine on its head. Once intubated, the tricaine concentration was reduced to 120 mg/ml while the fish and cloth were secured by a glass tube covering the chamber. The whole chamber was then inserted inside the probe, securely taped to avoid slippage and inserted into the 9.4T MRI scanner. Flowing intubating liquid was then switched from 120 mg/ml tricaine to 35mg/L of Benzocaine to ensure long-term safe sedation.

After scanning, fish were delicately taken off the chamber and deposited a tank of fresh E3 water at 23°C. Using a colder than usual temperature allowed the animal to adjust after being held in the scanner at ∼21 °C. Post scanning, fish recovery was recorded and animals were culled using overdose of tricaine and fixed in 4%PFA using the immersion-fixation regulated killing procedure.

### Gadolinium administration

Gadolinium MRI contrast agent (Gadovist→1.0 mmol/ml stock-Bayer Radiology, Leverkusen, Germany) was administered using two different routes:

1. For immersion, fish were left free swimming in a bath of 100 ml E3 containing 60 mmol/L of Gadovist (6.6 ml Gadovist in 100 ml of E3) for at least 2h.
2. For intra peritoneal (IP) injection: fish were anaesthetised using tricaine at 160 mg/ml until the fish stopped moving and responding to touch. Fish was transferred to a soft sponge laid in a petri dish and saturated with tricaine water at 160 mg/ml with a cut deep enough to accept the fish on its back. Using an insulin syringe (BD Micro-Fine + Demi U-100 Insulin 0.3 mm 30Gx8 mm, Fisher Scientific, Loughborough, UK), 5 μl of 1:2 Gadovist stock diluted in PBS (0.5 mmol/ml) was injected into the peritoneal cavity of the fish. The fish was then kept in 160 mg/ml tricaine to wrap in cloth and placed in the chamber.

### MRI Experiments

All scanning was performed on a 9.4T MRI (Bruker Biospec 70/30, Avance III spectrometer, 44 mm diameter vertical bore, 1500 mT/m gradient strength, Bruker Biospin MRI GmbH, Ettlingen, Germany) The probe was equipped with a 10 mm I.D. ^1^H/^13^C dual channel volume coil. Each animal was placed vertical into the probe with their head up. The chamber was positioned so that the fish’s head was located at the centre of a 10 mm 1H/13C volume rf coil (M2M, Cleveland, Ohio).

Scout images were acquired using FLASH (FOV 8×8 cm, matrix size 128×128, FA 30°, TE/TR 6/100 ms, 3 orthogonal slices, 2 mm slice thickness). To locate the brain more accurately two MSME spin echo scans were acquired in the axial and coronal planes (Axial FOV 1.2×0.8 cm; matrix size 80×60, TE/TR 8.5/1500 ms, 17 slices, 500 μm slice thickness. Coronal: FOV 1.2×0.8 cm; matrix size 128×64, TE/TR 14/1500 ms, 12 slices, 500 μm slice thickness:). Measurement of T1 and T2 was performed using a RARE-VTR sequence FOV 1.2×0.5 mm, 100×100 μm in plane resolution, 500 μm slice thickness, TE 10, 30, 50, 70 ms, TR 1000, 1282, 1674, 2333, 5000 ms). Further high-resolution scans (50×50 μm in plane resolution, 200 or 500 μm in plane resolution) were obtained with either MSME or RARE factor 2 scans, both TE/TR 14/1500ms. See figure legends for other details, e.g. NEX 16). Scan lasted between 45 to 90 minutes.

### Data Processing

Magnitude images were reconstructed without further modification using Bruker’s Paravision 5.1 software, these images were then imported into Matlab (R2018b, Mathworks, Natick, MA, USA) for further analysis. Brain regions of interest (ROI) were manually drawn by one of the authors (SR) for all image slices where brain tissue was observed. This region was further subdivided into fore, mid and hind regions, see supplementary data Figure S7. Each ROI was drawn on images extracted from the shortest TE, longest TR of the RARE-VTR data set. Subsequently, T1/T2 maps were calculated by fitting each voxel to the appropriate exponential equations (T1: Mo*(1-exp(-t/T1)+c; T2: Mo*(exp(-t/T1)+c) using the Matlab ‘Fit’ function. Any fitted T1 values > 7000 ms, T2 values > 300 ms or fit function derived r^2^ values < 0.98 were rejected. Statistical analysis was performed by one-way ANOVA and Bonferroni post-hoc test. Values are presented as mean ± standard error of mean (SEM) unless otherwise stated.

Signal to noise ratios were determined from voxels within the brain ROI for each slice. and a noise region within the slice. The images were same as used for defining ROIs from the RARE-VTR images. The noise region consisted of two locations in the bottom left and right portions from the head end of the image containing a voxel square covering 10% of the image along short axis. The SNR was calculated as 0.655*mean(ROI voxel intensity)/stdev(combined noise ROI)(Firbank et al., 1999).

## Supporting information

stepbystep protocol

## Notes

**Conflict of interest** The authors declare that the research was conducted in the absence of any commercial or financial relationships that could be construed as a potential conflict of interest.

### Competing Interest Statement

The authors have declared no competing interest.

