## Supplemental files for "Longitudinal brain studies in adult zebrafish by MRI"

**Figure S1**

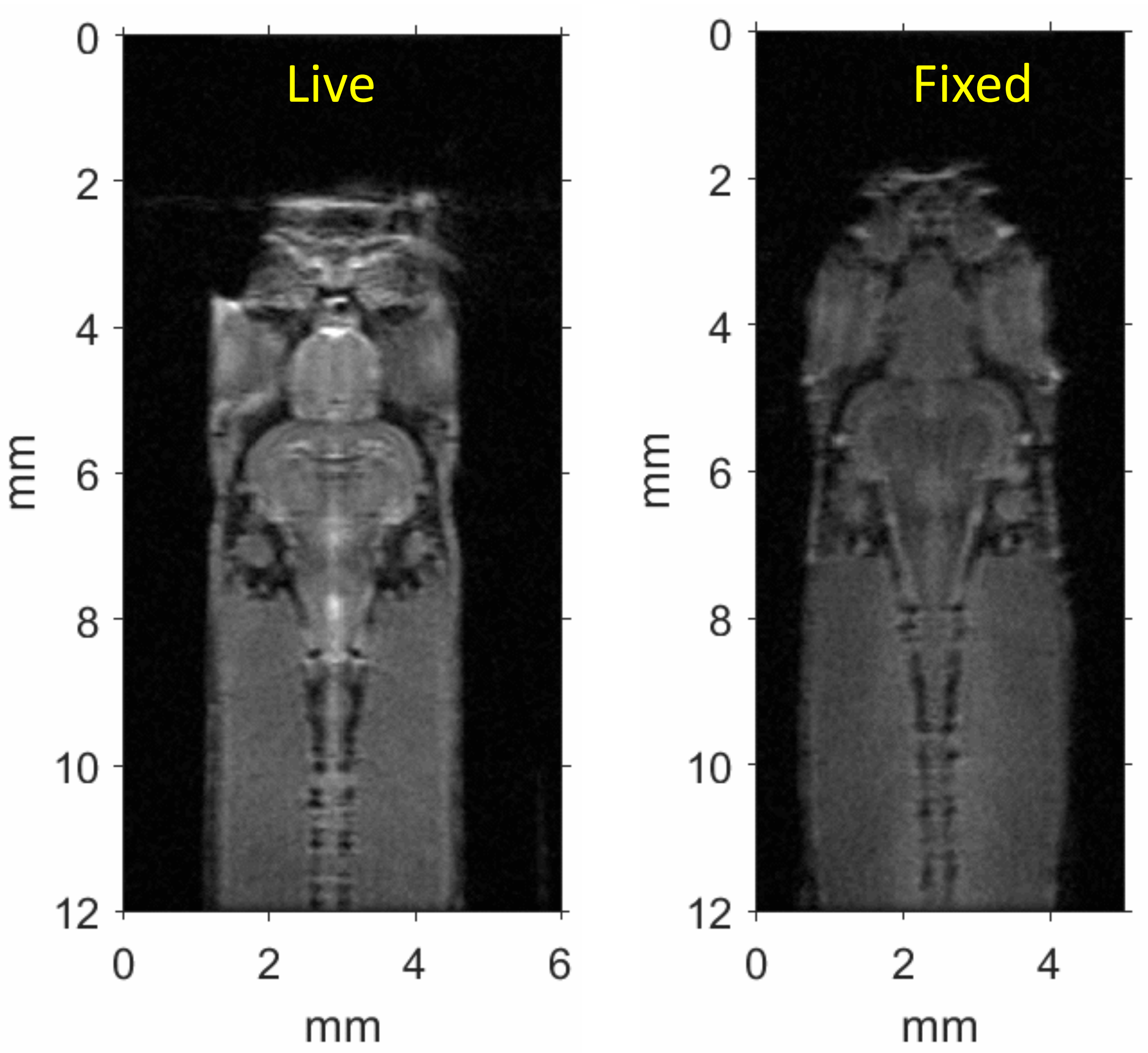


**Figure S1**: Representative RARE spin echo gif images from the same live (a) then fixed (b) zebrafish, with 50x50mm in plane resolution, 200mm slice thickness, TE/TR 14/1500ms, NEX 64. Live, FOV 12x6mm 96 min; Fixed, FOV 12x5mm, 80 min.

**Figure S2**

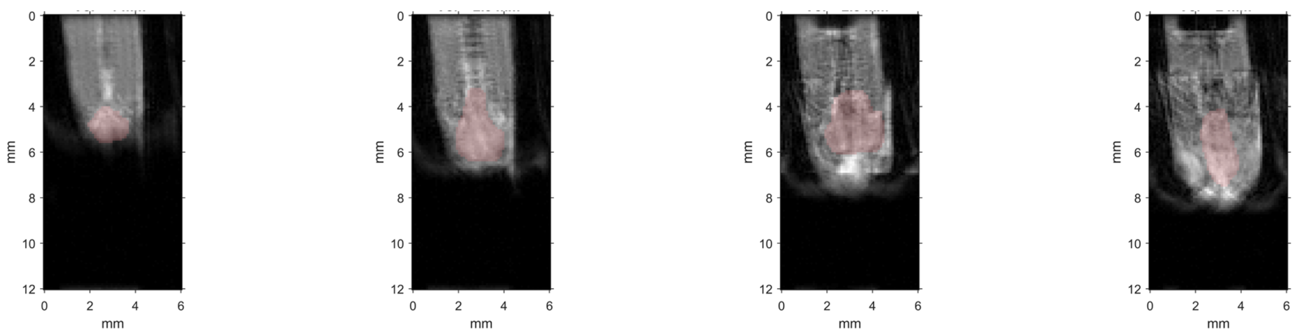


**Figure S2**: Motion artifacts observed in RARE spin echo image slicess from a live zebrafish (FOV 12x6mm, 50x50mm in plane resolution, 200mm slice thickness, TE/TR 14/1500ms, NEX 64, acquisition time 96 min).

### Figure S3

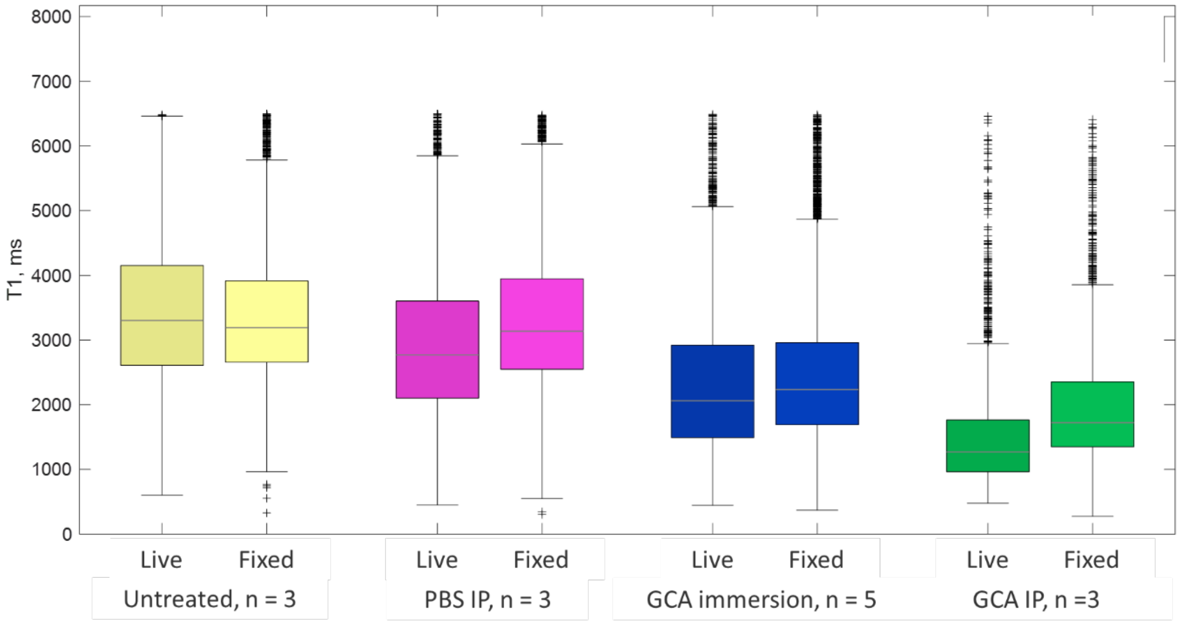


**Figure S3**: T1 measurement within the brain ROI of both live and fixed zebrafish. As for Figure 5A with outliers displayed and y-axis re-scaled. The boxplots show median and interquartile range (IQR), with whisker located at 1.5*IQR. See supplementary Tables S1 and S2 for live and fixed fish details respectively.

### Figure S4

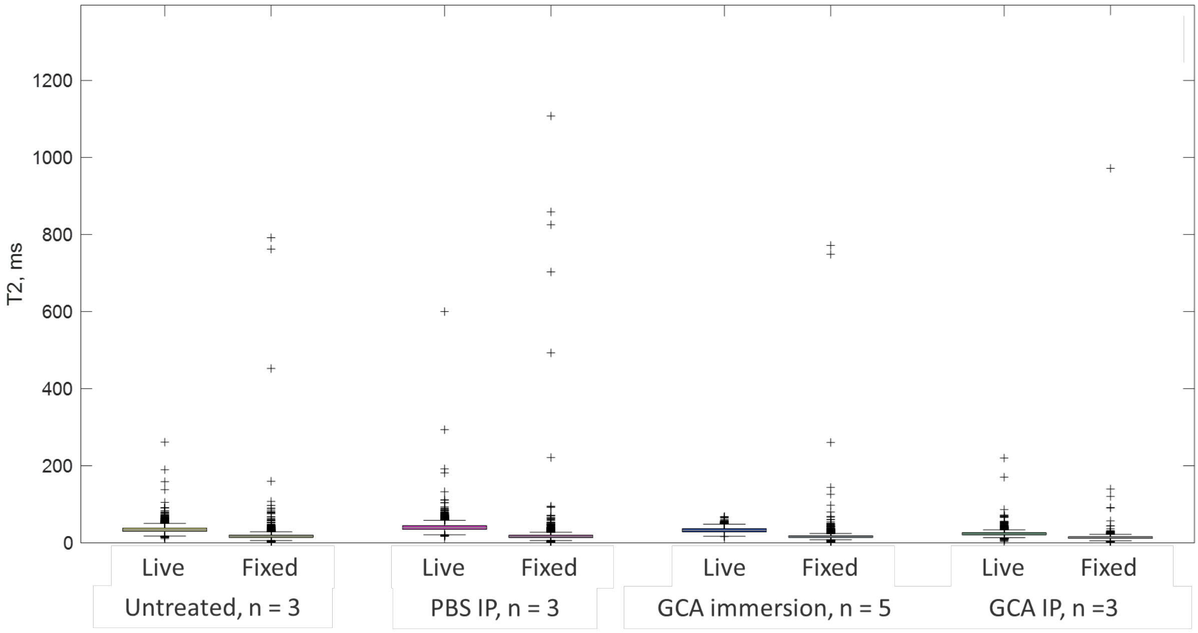


**Figure S4**: T2 measurement within the brain ROI of both live and fixed zebrafish. As for Figure 5B with outliers displayed and y-axis re-scaled. The boxplots show median and interquartile range (IQR), with whisker located at 1.5*IQR. See supplementary Tables S1 and S2 for live and fixed fish details respectively.

### Figure S5

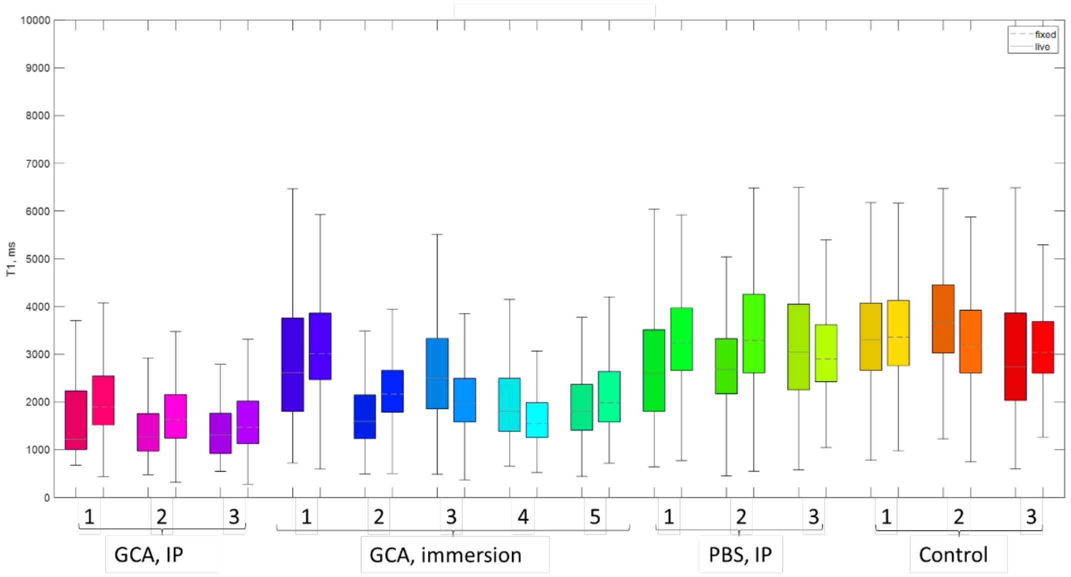


**Figure S5**: T1 value for live (left, solid median line in  box) and fixed (right, dashed line) for each fish. The boxplots show median and interquartile range (IQR), with whisker located at 1.5*IQR. Outliers have been removed for clarity.

### Figure S6

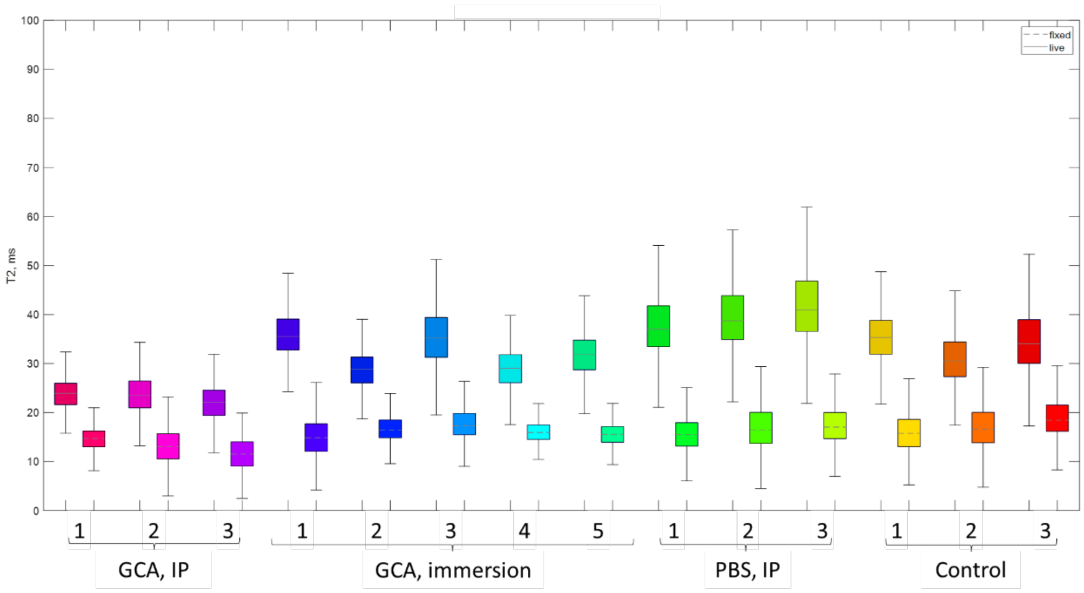


**Figure S6**: T2 value for live (left, with solid median line in  box) and fixed (right, with dashed line) for each fish. The boxplots show median and interquartile range (IQR), with whisker located at 1.5*IQR. Outliers have been removed for clarity.

**Figure S7**

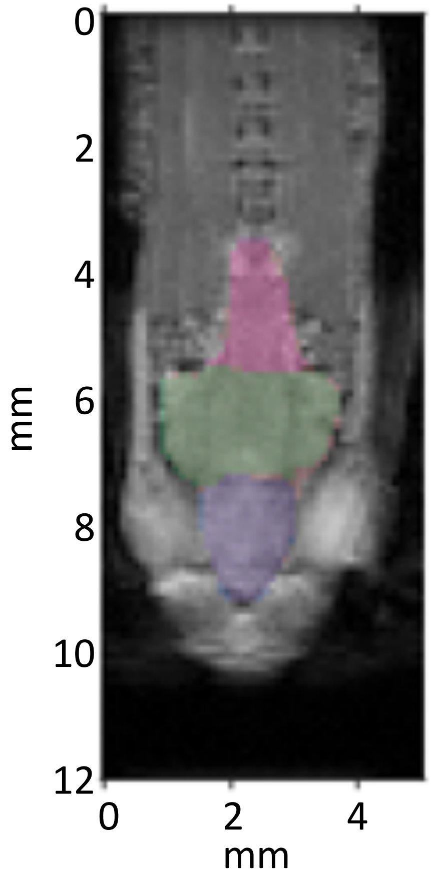


**Figure S7:** ROI regions manually drawn on an example RARE-VTR image (TE = 10 ms, TR 10000 s). Three ROI regions were defined; fore (approx. telencephalon, blue), mid (approx. optic tectum, green) and hind (approx. cerebellum and part of brain stem, red) in each slice, where the brain region was visible.

### Figure S8

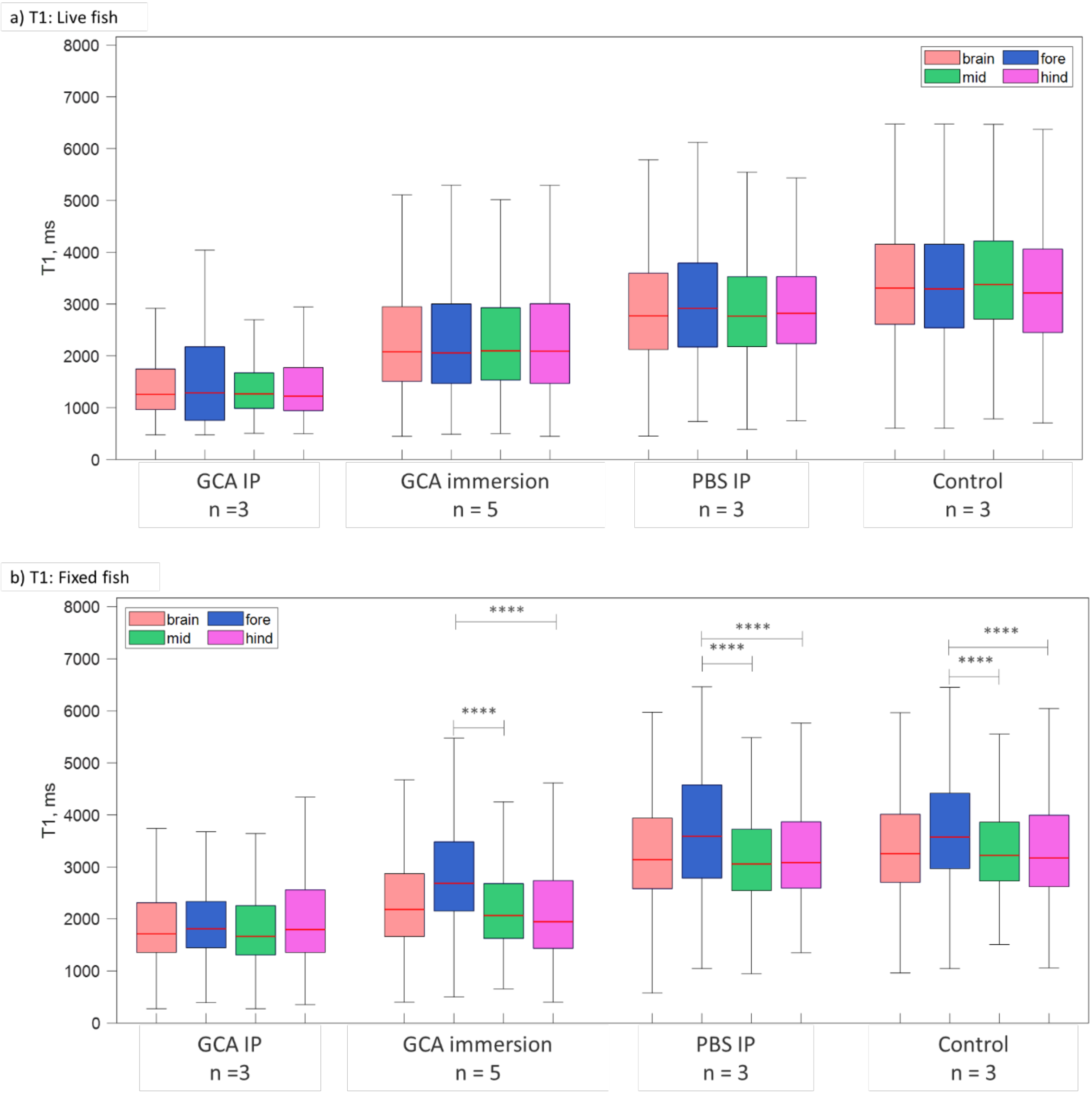


**Figure S8:** T1 values within differing regions of interest of the brain (fore, mid and hind, see Figure S1) for both live (a) and fixed (b) zebrafish. Prior to scanning, live fish were treated with gadolinium contrast agent (GCA) either, IP injected or immersion in a solution of it. Controls received either an IP injection of PBS or immersion in standard fish tank water (see main text for details). The boxplots show median and interquartile range (IQR), with whisker located at 1.5*IQR. Outliers have been removed for clarity. The T1 value from the whole brain region are the same as Figure 4 and are included as a guide. Horizontal bars show Bonferroni Post-hoc differences between fixed fish cohorts (there were no significant differences for the live fish). See supplementary Table S4 for fixed fish details. ****, $p << 0.0001$.

### Figure S9

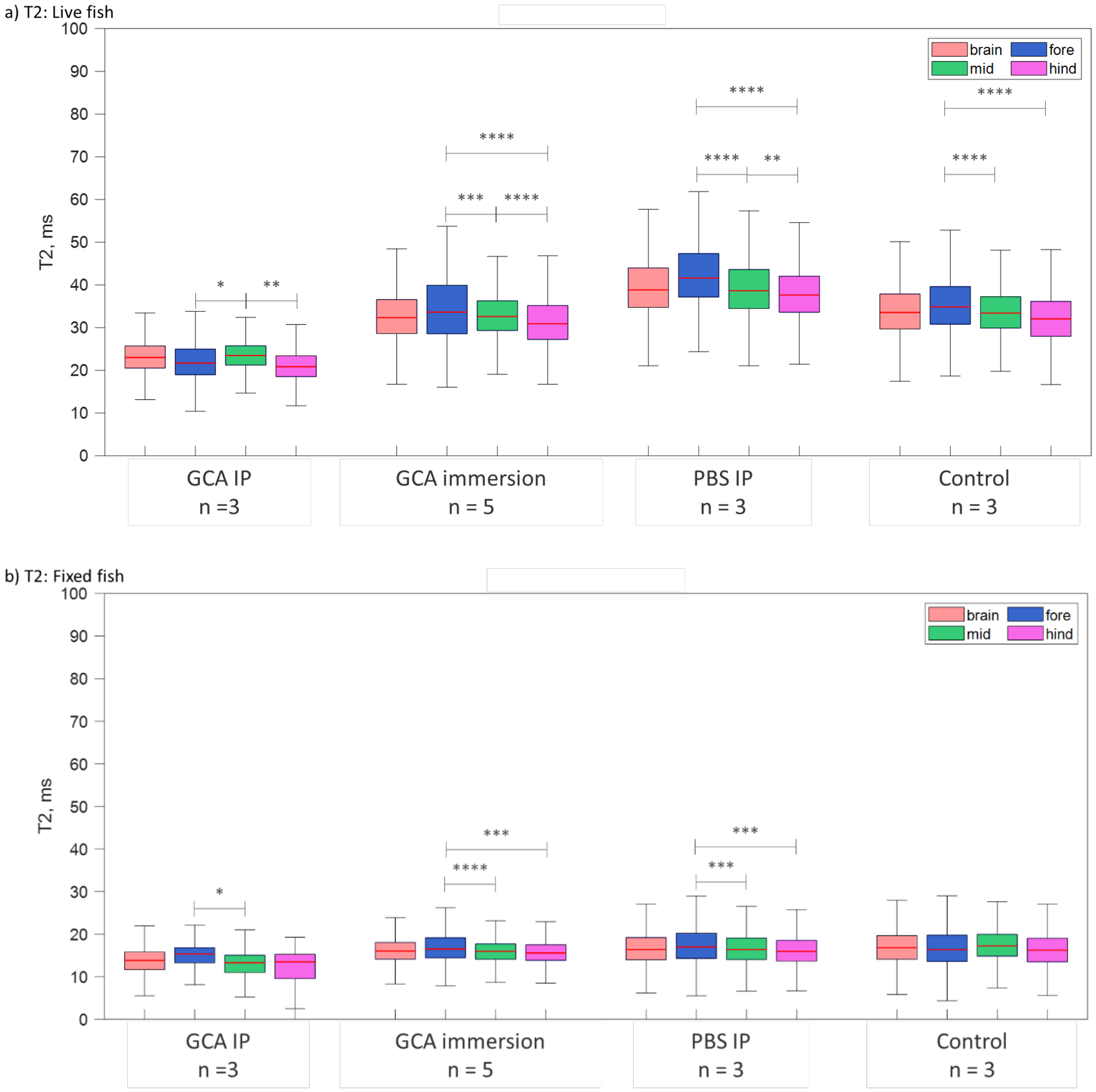


**Figure S9:** T2 values within differing regions of interest of the brain (fore, mid and hind, see Figure S1) for both live (a) and fixed (b) zebrafish. Prior to scanning, live fish were treated with gadolinium contrast agent (GCA) either, IP injected or immersion in a solution of it. Controls received either an IP injection of PBS or immersion in standard fish tank water (see main text for details). The boxplots show median and interquartile range (IQR), with whisker located at 1.5*IQR. Outliers have been removed for clarity. Horizontal bars show Bonferroni Post-hoc differences between fish cohorts. The T2 value from the whole brain region are the same as Figure 5 and are included as a guide. See supplementary Tables S5 and S6 for live and fixed fish details respectively. * $p < 0.05$, ** $p < 0.01$, *** $p < 0.001$, ****, $p << 0.0001$.

**Table S1**: Comparison of T1 values between live or fixed fish. P-values were determined by a Bonferroni Post-hoc applied to a one-way ANOVA (IP, intra peritoneal injection; immer, GCA administered by immersion).

| **T1, ms** | Group 1 | Mean ± SEM | Group 2 | Mean ± SEM | p-value |
|---|---|---|---|---|---|
| Live | GCA IP | 1590 ± 27 | GCA immer | 2341 ± 20 | << 0.0001 |
| | GCA IP | 1590 ± 27 | PBS IP | 2955 ± 20 | << 0.0001 |
| | GCA IP | 1590 ± 27 | Control | 3426 ± 19 | << 0.0001 |
| | GCA immer | 2341 ± 20 | PBS IP | 2955 ± 20 | << 0.0001 |
| | GCA immer | 2341 ± 20 | Control | 3426 ± 19 | << 0.0001 |
| | PBS IP | 2955 ± 20 | Control | 3426 ± 19 | << 0.0001 |
| Fixed | GCA IP | 1971 ± 20 | GCA immer | 2466 ± 13 | << 0.0001 |
| | GCA IP | 1971 ± 20 | PBS IP | 3334 ± 16 | << 0.0001 |
| | GCA IP | 1971 ± 20 | Control | 3374 ± 15 | << 0.0001 |
| | GCA immer | 2466 ± 13 | PBS IP | 3334 ± 16 | << 0.0001 |
| | GCA immer | 2466 ± 13 | Control | 3374 ± 15 | << 0.0001 |

**Table S2**: Comparison of T2 values between live or fixed fish. P-values were determined by a Bonferroni Post-hoc applied to a one-way ANOVA (IP, intra peritoneal injection; immer, GCA administered by immersion).

| **T2, ms** | Group 1 | Mean ± SEM | Group 2 | Mean ± SEM | p-value |
|---|---|---|---|---|---|
| Live | GCA IP | 24.3 ± 0.2 | GCA immer | 32.8 ± 0.1 | << 0.0001 |
| | GCA IP | 24.3 ± 0.2 | PBS IP | 40.4 ± 0.2 | << 0.0001 |
| | GCA IP | 24.3 ± 0.2 | Control | 34.4 ± 0.1 | << 0.0001 |
| | GCA immer | 32.8 ± 0.1 | PBS IP | 40.4 ± 0.2 | << 0.0001 |
| | GCA immer | 32.8 ± 0.1 | Control | 34.4 ± 0.1 | << 0.0001 |
| | PBS IP | 40.4 ± 0.2 | Control | 34.4 ± 0.1 | << 0.0001 |
| Fixed | GCA IP | 14.1 ± 0.3 | GCA immer | 16.5 ± 0.1 | << 0.0001 |
| | GCA IP | 14.1 ± 0.3 | PBS IP | 17.8 ± 0.3 | << 0.0001 |
| | GCA IP | 14.1 ± 0.3 | Control | 18.0 ± 0.2 | << 0.0001 |
| | GCA immer | 16.5 ± 0.1 | PBS IP | 17.8 ± 0.3 | << 0.0001 |
| | GCA immer | 16.5 ± 0.1 | Control | 18.0 ± 0.2 | << 0.0001 |

**Table S3**: Bonferroni derived p-value for comparison of T1 or T2 values between Live vs Fixed fish. n.s. not significant. (IP, intra peritoneal injection; immer, GCA administered by immersion).

| Live vs Fixed | T1 p-value | T2 p-value |
|---|---|---|
| GCA IP | << 0.0001 | << 0.0001 |
| GCA immer | << 0.0001 | << 0.0001 |
| PBS IP | << 0.0001 | << 0.0001 |
| Control | n.s. | << 0.0001 |

**Table S4**:  T1 values between different brain ROIs (fore, mid, hind, see Figure 3) in fixed fish. Values were compared using a one-way ANOVA with between groups p-values determined by a Bonferroni Post-hoc test (IP, intra peritoneal injection; immer, GCA administered by immersion).

| **Fixed fish. T1, ms** | | | | | |
|---|---|---|---|---|---|
| Group 1 | ROI 1 | Mean ± SEM | ROI 2 | Mean ± SEM | p-value |
| GCA immer | fore | 2946 ± 28 | mid | 2280 ± 16 | << 0.0001 |
| GCA immer | fore | 2946 ± 28 | hind | 2187 ± 31 | << 0.0001 |
| PBS IP | fore | 3722 ± 42 | mid | 3243 ± 21 | << 0.0001 |
| PBS IP | fore | 3722 ± 42 | hind | 3316 ± 33 | << 0.0001 |
| Control | fore | 3720 ± 37 | mid | 3393 ± 32 | << 0.0001 |
| Fixed: GCA IP | fore | 3720 ± 37 | hind | 3412 ± 32 | << 0.0001 |

**Table S5**: T2 values between different brain ROIs (fore, mid, hind, see Figure 3) in live fish. Values were compared using a one-way ANOVA with between groups p-values determined by a Bonferroni Post-hoc test (IP, intra peritoneal injection; immer, GCA administered by immersion).

| **Live fish. T2, ms** | | | | | |
|---|---|---|---|---|---|
| Group 1 | ROI 1 | Mean ± SEM | ROI 2 | Mean ± SEM | p-value |
| GCA IP | mid | 24.0 ± 0.1 | fore | 22.2 ± 0.2 | 0.03 |
| GCA IP | mid | 24.0 ± 0.1 | hind | 21.9 ± 0.4 | < 0.01 |
| GCA immer | fore | 34.3 ± 0.3 | mid | 33.0 ± 0.1 | < 0.001 |
| GCA immer | fore | 34.3 ± 0.3 | hind | 31.6 ± 0.2 | << 0.0001 |
| GCA immer | mid | 33.0 ± 0.1 | hind | 31.6 ± 0.2 | << 0.0001 |
| PBS IP | fore | 43.2 ± 0.3 | mid | 39.7 ± 0.2 | << 0.0001 |
| PBS IP | fore | 43.2 ± 0.3 | hind | 38.3 ± 0.3 | << 0.0001 |
| PBS IP | mid | 39.7 ± 0.2 | hind | 38.3 ± 0.3 | < 0.01 |
| Control | fore | 35.6 ± 0.2 | mid | 34.0 ± 0.1 | << 0.0001 |
| Control | fore | 35.6 ± 0.2 | hind | 33.0 ± 0.4 | << 0.0001 |

**Table S6**: T2 values between different brain ROIs (fore, mid, hind, see Figure 3) in fixed fish. Values were compared using a one-way ANOVA with between groups p-values determined by a Bonferroni Post-hoc test (IP, intra peritoneal injection; immer, GCA administered by immersion).

| **Fixed fish. T2, ms** | | | | | |
|---|---|---|---|---|---|
| Group 1 | ROI 1 | Mean ± SEM | ROI 2 | Mean ± SEM | p-value |
| GCA IP | fore | 15.2 ± 0.1 | mid | 13.2 ± 0.1 | 0.04 |
| GCA immer | fore | 17.9 ± 0.6 | mid | 16.0 ± 0.1 | < 0.0001 |
| GCA immer | fore | 17.9 ± 0.6 | hind | 15.8 ± 0.1 | < 0.001 |
| PBS IP | fore | 18.9 ± 0.6 | mid | 16.8 ± 0.1 | < 0.001 |
| PBS IP | fore | 18.9 ± 0.6 | hind | 16.6 ± 0.1 | < 0.001 |
