## Supplementary material for "Longitudinal brain studies in adult zebrafish by MRI": stepbystep protocol

### Step-by-Step protocol

#### 1. Reagents, software and equipment:

#### 1.1. For anaesthesia, Gadolinium administration and fixing

- E3: 10x E3 (5mM NaCl, 0.17mM KCl, 0.33mM CaCl2, 0.33mM MgSO4) fish water to dilute to 1x E3 with distilled water (Kimmel et al. 1995).
- Tricaine/MS222 (Sigma – Cat. No. E10521, Gillingham, UK) stock was made at 4g/L in filtered system water and bring to pH 7.4 using 1M Tris. Stock was stored in the fridge in a brown glass bottle to protect from light. Subsequent tricaine solutions were made using Tricaine stock and E3 water: 160mg/ml of tricaine (4ml in 96ml of E3), 120mg/ml tricaine (3ml in 96ml of E3).
- Benzocaine (Sigma E1501) stock at 50mg/ml was made by dissolving in 70% EtOH, aliquots of 750ul were stored at -20 until usage. Benzocaine was used at the final concentration of 35mg/L (using 700ul in 1L to feed the pumps).
- Multipurpose cloth (Spontex CODE/SKU: 19900074)
- Gadovist®1.0 mmol/ml stock- Bayer Radiology, Leverkusen, Germany
- Soft sponge used for washing up (Jantex sponge)
- Syringes for IP injection insulin syringe (BD Micro-Fine + Demi U-100 Insulin 0.3 mm 30Gx8 mm, Fisher Scientific, Loughborough, UK)
- PARAFILM®M

#### 1.2 For water supply and scanning

- 9.4T MRI (Bruker Biospec 70/30, Avance III spectrometer, 44 mm diameter vertical bore, 1500 mT/m gradient strength, Bruker Biospin MRI GmbH, Ettlingen, Germany)
- Probe with 10 mm I.D. $^{1}H/^{13}C$ dual channel volume coil (M2M, Cleveland, Ohio)
- Two Aladdin NE-1000 syringe pumps (World Precision Instruments, Hitchin, UK)
- Syringes for pumps (Terumo Syringe sterile without needle 10ml SS+10ES1)

#### 1.3 For chamber 3D design and printing:

- Polylactic acid for 3D printing
- Open-source software Blender (www.blender.org)
- Ultimaker 2+ Extended 3D printer with 3D printed using Cura software (both Ultimaker, Utrecht, Netherlands)
- Acrylic rods (1 mm diameter, ~5 cm in length. 4D Modelshop Ltd, London, UK)
- ~20 cm, 0.25 mm I.D PVC tube extended with a 0.89 mm I.D tube approximately 2 m in length (S3 and S54-HL Tygon respectively, Cole Parmer, St Neots, UK).
- 10 mm, flat bottom, glass NMR tube (DWK, Stoke on Trent, UK)
- 8 x 2 mm o-rings, (RS components, Corby, UK)

#### 2. Methodology:

The day before scanning, defrost 4% PFA and store at 4°C to use for fish fixing post-scan.
A day of scanning consists of 4 animals scanned back-to-back. Animals were not fed on the morning of the scan and travelled together in a travel tank placed in a polystyrene box from the aquarium to the MRI facility. Fish were stored in a travel tank placed in a polystyrene box to reduce heat loss during the day of scanning.

#### 2.1 Preparation of anaesthetic solutions

-Tricaine/MS222 (Sigma – Cat. No. E10521, Gillingham, UK) stock was made at 4 g/L in filtered system water, with pH adjusted to 7.4 using 1 M Tris. A brown glass bottle protected the stock solution from light and was stored in the fridge. Subsequent tricaine solutions were made using Tricaine stock and E3 water (10x E3: 5 mM NaCl, 0.17 mM KCl, 0.33 mM $CaCl_2$, 0.33 mM $MgSO_4$ diluted to 1x E3 with distilled water (Kimmel et al. 1995). The anaesthetic solution concentrations were made with high accuracy: 160 mg/ml of tricaine (4ml in 96ml of E3) and 120 mg/ml tricaine (3 ml in 96 ml of E3).
-Benzocaine (Sigma E1501) stock was made at 50 mg/ml by dissolving in 70% ethanol. This was aliquoted at 750 ml and stored at -20°C until usage. Benzocaine was used at the final concentration of 35 mg/L (using 700 ml in 1 L to directly feed the syringe pumps) and made fresh at the start of each scanning day.

**2.2 Immersion in Gadolinium (optional)**
To give enough time for the gadolinium immersion group to swim in the contrast agent, we placed one fish in a glass beaker with 100ml of E3 containing 60mmol/L of Gadovist (6.6ml in 100ml) for at least 2h of free swimming. The tank was placed in a polystyrene box to conserve the temperature of the water. During immersion time, we started to prepare our untreated control fish of the day.

**2.3 Intra peritoneal (IP) Gadolinium injection (optional)**
**Prepare the surgical table/sponge:** Make a deep cut in a soft sponge (such as #L800-D, Jaece Industries) to use as a holding trough to hold the fish during injection. Saturate the sponge with E3 containing 160 mg/ml tricaine and place in a dish containing more tricaine (we used a P200 pipette tip box lid). Set the sponge into a 60 mm Petri dish. Set the Petri dish with sponge into a suitably-sized pipette tip box lid. The lid needs to be large enough to hold water to help maintain the sponge moist, but it should be shallow enough to not get in the way.
**Prepare syringe:** a 5ml drop of 1:2 Gadovist stock diluted in PBS (0.5 mmol/ml) was placed on a clean piece of parafilm and taken up using an insulin syringe (BD Micro-Fine + Demi U-100 Insulin 0.3 mm 30Gx8 mm, Fisher Scientific, Loughborough, UK), ensuring no air bubble entered the syringe.
**Anaesthesia and injection**: Fish were anaesthetised using tricaine at 160 mg/ml until the fish stopped moving and responding to touch. Fish was transferred to the prepared slit in the sponge belly up and carefully insert the needle into the midline between the pelvic fins. The needle should point cranially and be inserted closer to the pelvic girdle than to the anus. You should be able to feel when the needle is deep to the body wall. Inject the appropriate volume and withdraw the needle. This JoVe video contains all the information necessary (Kinkel et al. 2010) https://www.jove.com/v/2126/intraperitoneal-injection-into-adult-zebrafish. The fish was then taken out of the sponge and placed back in a petri dish containing 160 mg/ml tricaine for positioning into the chamber (see next steps below).

**2.4 Zebrafish anaesthesia, scanning and recovery**
For initial anaesthesia, fish were transferred from their holding tank using a net to a beaker containing 100 ml of 160 mg/ml tricaine. When the fish stopped moving and responding to touch, it was lifted out of the beaker and transferred to a shallow bath of 160 mg/ml tricaine in a clean petri dish. There, the fish was wrapped into a thin rectangle of multipurpose cloth (Spontex CODE/SKU: 19900074), which allowed the fish to be gently held vertically without slipping or damaging its scales (Figure 1A). Fish were vertically inserted between the pliable rods of the live chamber (Figure 1B), ensuring the feeding tube was positioned ventrally to stop any water artefacts appearing on the side of the brain. During the positioning process, the fish was kept moist by dripping drops of 160 mg/ml tricaine on its head. Once intubated, the tricaine concentration was reduced to 120 mg/ml while the fish and cloth (Figure 1C) were secured by a glass tube covering the chamber.

The whole chamber was then inserted inside the MRI probe, securely taped to avoid slippage and inserted into the 9.4T MRI scanner (Figure 1D). Flowing intubating liquid was then switched from 120 mg/ml tricaine to 35mg/L of Benzocaine to ensure long-term safe sedation for the rest of the experiment.

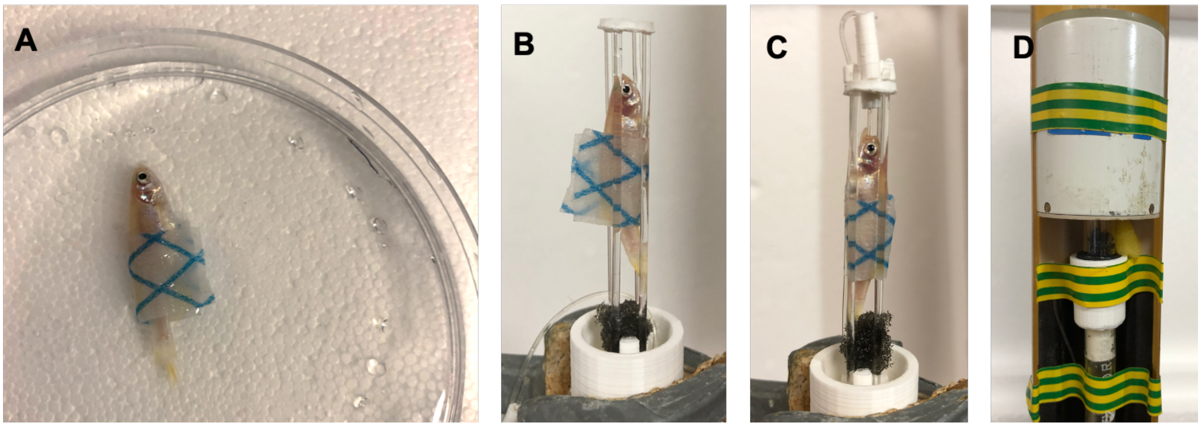


**Figure 1**: Series of images taken during the setup of the fish from initial anaesthesia (A), to positioning (B) and intubating (C), and finally insertion of the fish and probe inside the scanner (D).

After scanning, fish were delicately taken off the chamber and deposited in a tank of fresh E3 water at 23°C. Using a colder than usual temperature allowed the animal to adjust after being held in the scanner at ~21 °C. We recorded the recovery of each scanned fish for future longitudinal studies.
Fish were then culled using regulated immerse/fixation killing: fish were transferred with a net to a beaker containing lethal dose of Tricaine MS-222 and transferred to cold 4% PFA as soon as the fish stopped reacting to movement. Fixed fish were kept in the fridge for 2 days before being mounted in a new chamber (to avoid PFA contamination) for measuring T1 and T2 signals of fixed tissues.

### Additional references


Kimmel, Charles B., William W. Ballard, Seth R. Kimmel, Bonnie Ullmann, and Thomas F. Schilling. 1995. *Stages of Embryonic Development of the Zebrafish*.

Kinkel, Mary D., Stefani C. Eames, Louis H. Philipson, and Victoria E. Prince. 2010. “Intraperitoneal Injection into Adult Zebrafish.” *Journal of Visualized Experiments* (42). doi: 10.3791/2126.
